# Population Resilience Under Environmental Deterioration in Socially Monogamous Systems with Mutual Mate Choice

**DOI:** 10.64898/2026.06.01.729368

**Authors:** Neelam Porwal, Jonathan M. Parrett, Franky Rogers, Jacek Radwan, Robert J. Knell

**Affiliations:** Evolutionary Biology Group, Faculty of Biology, Adam Mickiewicz University, Poznań, Poland; School of Environmental Sciences, University of East Anglia, Norwich, England, Great Britain; School of Environmental and Life Sciences, University of Hull, Hull, United Kingdom

**Author notes:** **Corresponding authors:** Robert J. Knell, Neelam Porwal. **Author Contributions:** NP and RJK conceived the ideas and designed the methodology with the support of FR, JR, and JMP; NP wrote the code with support from FR and RJK. NP, RJK, JR, and JMP interpreted the results. NP led the writing of the manuscript with contributions from RJK, JR, and JMP. All authors gave final approval for publication. **Competing Interest Statement:** The authors declare no competing interests.

**Keywords:** Extra-pair copulations, Extra-pair paternity, Extinction risk, Population resilience, Small populations

## Abstract

Rapid environmental change and biodiversity loss make it increasingly important to identify factors influencing population extinction risk. Previous studies examining how mating systems can affect persistence of populations under environmental stress generally report higher extinction risks in monogamous than polygynous systems but have largely ignored extra-pair copulations (EPC) and paternity (EPP), despite the prevalence of genetic polyandry in socially monogamous species. Here, using an individual-based model, we study how EPP in socially monogamous systems affects population resilience under directional environmental change. We assume that in socially monogamous species, both sexes carry costly sexual ornaments, the elaboration of which depends on the strength of preference. The effect of EPPs on extinction risk depended on the strength of mate preference, population size, and the degree to which homozygosity affected fitness. Systems with EPCs are not simply intermediate in resilience between strict monogamy and polygyny: the preference strength interacts with mating system, leading to superior resilience of EPC systems compared to strictly monogamous and polygynous systems when choosiness and the negative consequences of heterozygosity loss are low, and EPP rates are high. However, this benefit was reduced in small populations due to faster loss of heterozygosity. At high choosiness, EPC systems exhibited lower resilience than socially polygynous choice systems because the higher reproductive skew of the latter system allowed them to adapt faster while not suffering from the demographic consequences of sexual signaling costs borne by females. Overall, our results suggest that EPCs can enhance population resilience when females obtain fertilizations from higher-condition extra-pair males compared to systems without EPC.

## Introduction

Present-day extinction rates have escalated to unprecedented levels due to intensifying anthropogenic disturbances, especially those associated with rapid climate change (Román-Palacios & Wiens, 2020; Urban, 2024). These accelerating environmental shifts raise an evolutionary question: whether species can adapt rapidly enough to keep pace with ongoing climatic change (Lavergne et al., 2010; Salamin et al., 2010), making understanding how populations adapt to environmental change an important goal for contemporary biologists. The strength of sexual selection is known to influence population dynamics, recovery potential, and extinction risk, with empirical, theoretical, and comparative studies documenting positive (Godwin et al., 2020; Lumley et al., 2015; Martínez-Ruiz & Knell, 2017; Martinossi-Allibert et al., 2018; Moore et al., 2024; Parrett et al., 2019), negative (Bro-Jørgensen, 2014; Martínez-Ruiz & Knell, 2017; Martins et al., 2018; Sorci et al., 1998), and neutral (Morrow & Fricke, 2004; Morrow & Pitcher, 2003) effects across diverse taxa. This inconsistency between the results may be a consequence of sexual selection being modulated via the mating strategies, systems, and mate choice modes in which it is occurring. However, while a substantial body of research has examined how rapid environmental change affects population persistence in the context of strong or weak sexual selection, far less attention has been devoted to understanding how mating system dynamics shape population resilience in these contexts (but see Porwal et al 2026).

Experimental studies comparing polygamous and monogamous systems under stress have typically enforced monogamy, thereby eliminating opportunities for mate choice to operate. Under these conditions, monogamous systems have often performed worse than polygamous systems (Godwin et al., 2020; Lumley et al., 2015; Martinossi-Allibert et al., 2018), and a generally positive trend was inferred from meta-analyses of these studies (Cally et al., 2019). Previous theoretical models comparing mating systems have generally implied greater extinction risks in monogamous populations relative to polygamous ones (Leach et al., 2020; Legendre et al., 1999; Sæther et al., 2004). However, extinction risks are contingent on sex ratios in the populations and can also produce opposite outcomes (Bessa-Gomes et al., 2004; Lee et al., 2011). These studies assessed extinction risk primarily through stochastic processes in stable environments. Furthermore, they did not account for the effects of heterozygosity loss or explicitly model mate choice direction and strength. (Leach et al., 2020; Legendre et al., 1999; Sæther et al., 2004; Bessa-Gomes et al., 2004). More recently, Porwal et al. (2026) modelled a variety of mating systems and compared population resilience under environmental change, finding that monogamous systems with mutual choice exhibited lower resilience when compared with polygynous systems with female choice. For simplicity, however, this study modelled strict genetic monogamy, which, although present in some species, is not typical of many socially monogamous systems, which often feature some degree of extra-pair copulation (EPC) and extra-pair paternity (EPP) (Cockburn, 2006; Hasselquist & Sherman, 2001).

In socially monogamous birds with biparental care, for example, genetic polyandry has been documented in more than 70% of species. On average, approximately 19% of offspring in these species are sired by extra-pair males, with extra-pair paternity (EPP) occurring in around 33% of broods (Brouwer & Griffith, 2019). The highest occurrence of EPP (88.9% broods with EPP, Whittingham & Dunn, 2001) in socially monogamous species, populations are reported from Tree Swallows, *Tachycineta bicolor*, with 68.5% of offspring sired by extra-pair males (Barber et al., 1996). This is followed by the New Zealand Hihi, *Notiomystis cincta*, with a reported EPP of 68% occurring in 88.9% of the broods (Brekke et al., 2013). Considerable interspecific variation exists in both EPC frequency and rates of extra-pair paternity, leading to the development of multiple hypotheses to explain this diversity (Wan et al., 2013). Among these, the adaptive benefits hypothesis, particularly the indirect genetic benefits gained by females, remains one of the most widely supported explanations. Apart from the individual benefits of trading up for a “better” male for females, males benefit by increasing their reproductive success (Kempenaers & Dhondt, 1993) extra-pair matings may also facilitate the spread of beneficial alleles within populations (Akçay & Roughgarden, 2007; Brouwer & Griffith, 2019; Forstmeier et al., 2014; Wan et al., 2013). Under the adaptive benefits hypothesis, the “good genes” hypothesis proposes that females paired with lower-quality males may engage in EPCs to obtain fertilizations from genetically superior males (Akçay & Roughgarden, 2007; Arct et al., 2015; Brouwer & Griffith, 2019; Griffith et al., 2002; Jennions & Petrie, 2000; Johnsen et al., 1998; Kempenaers et al., 1997; Otter et al., 1998; Petrie & Kempenaers, 1998; Rätti et al., 1995; Saino et al., 1997; Strohbach et al., 1998; Wan et al., 2013; Zeh & Zeh, 2001). Such genetic benefits may become especially important under rapidly changing environmental conditions and could influence extinction risk in socially monogamous systems.

Biparental care is common in socially monogamous systems, and it often selects for choosiness in both sexes (Jones & Hunter, 1993; Kokko & Johnstone, 2002). Under these conditions, both sexes may evolve preferences for traits that reliably signal individual condition and quality (Baldauf et al., 2009; Beeching & Hopp, 1999; Holveck & Riebel, 2010; Jiang et al., 2013; Kirkpatrick et al., 2006; Kraak & Bakker, 1998; McKaye, 1986; Rueger et al., 2016). Such reliable signalling requires elaborate and energetically costly traits in both sexes (Aquiloni & Gherardi, 2008; Holveck & Riebel, 2010). The production and maintenance of costly sexual traits may also increase mortality risks for both males and females (Hooper & Miller, 2008). The strength of choosiness/preference, translating into the strength of assortative mating, may therefore have important evolutionary consequences, influencing both the maintenance of genetic variation and the direction of selection within populations (Bolnick & Kirkpatrick, 2012). Hence, the strength of choosiness may further complicate evolutionary responses of mutual choice in socially monogamous systems under environmental change.

Here, we extend the framework of the model by Porwal et al. (2026) by incorporating EPCs.

Assuming that females engage in EPCs to gain indirect benefits under the good gene hypothesis, we vary the strength of choosiness and investigate how the rate of extra-pair paternity when EPC occurs affects population resilience to adverse environmental change.

## Methods

The individual-based simulation model used here extends the framework developed by Porwal et al. (2026) to investigate how extra-pair copulations (EPCs) influence evolutionary dynamics under environmental change in socially monogamous systems with mutual mate choice. To benchmark the effects of EPCs, we compared mutual-choice monogamy with and without EPCs to polygynous systems with and without mate choice under varying levels of the strength of choosiness. The model retained the mixed genetic architecture, condition-dependent signaling, heterozygosity dynamics, density dependence, and environmental change processes described in Porwal et al. (2026). Simulations were implemented in R version 4.5.2 (R Core Team, 2023) as a single simulation function following Acerbi et al. (2022).

As in Porwal et al. (2026), genome-wide heterozygosity was calculated by comparing 50 neutral loci on two strands where these loci differed. The environment was modelled as a single continuous variable, and phenotype, in this case, the optimal value of environment for an individual, was modelled as a continuous trait based on Fisher’s infinitesimal model. Subsequently, individual condition depended on environmental mismatch (the absolute difference between the environment and individual phenotype) and heterozygosity, with mismatch and homozygosity penalties reducing signaling expression, survival, and reproductive success. Environmental conditions fluctuated stochastically or changed directionally following an initial 50-time-step stabilization period. Simulations ended either after 300 time steps or at extinction, defined as the loss of all adults and immatures (Brook et al., 2008).

Mating followed the group-based ranking procedure described by Porwal (2026), adapted from (Petrie et al., 1991). Females were assigned to mating groups of up to ten individuals, after which males were distributed randomly among groups. In mutual-choice systems, both sexes were ranked according to signaling trait expression, and mating probabilities were determined by the mate preference strength (*β*). Monogamous systems enforced one-to-one pairing by removing unpaired lowest-ranking males or females (depending on which sex was more numerous within a group) (Bessa-Gomes et al., 2004; Legendre et al., 1999), reflecting the fact that random mortality may lead to an excess of adults of one or the other sex (Haridas et al., 2014; Kvarnemo et al., 2006; Sun et al., 2022). Whereas polygynous systems allowed males to mate with multiple females, resulting in mating with even lower quality females (Kvarnemo, 2018). Unlike in Porwal et al. 2026 where β was fixed, here we varied it to explore how preference strength influences population outcomes.

To include EPCs in mutual-choice monogamy systems, after pairings were determined, females could obtain extra-pair fertilizations from males with higher signaling values than their social mate, consistent with the “good genes” hypotheses (Akçay & Roughgarden, 2007; Wan et al., 2013). After pairing up in the first round, the benefits for males to engage in further matings lie in improving their reproductive success without having to provide parental care (Kampenaers & Dhont, 1993). Hence, they did not have a preference for females, so while males were choosy for their social partner, they were not choosy for EPC partners; females expressed preference for both social and EPC partners. EPC probability increased with the signaling difference between the female’s social mate and the highest-ranked male in the mating group (Hasselquist et al., 1996; Kempenaers et al., 1992; Landgraf et al., 2017; Mennill et al., 2003; Møller & Birkhead, 1994; Otter et al., 1998; Whittingham & Dunn, 2016) and with the predefined EPC parameter determining the likelihood of EPC occurrence. When EPCs occurred, fertilization was assigned probabilistically in the respective groups among males ranking higher than the social male according to male rank (based on the signal trait) and *β*.

Reproductive output in mutual-choice monogamy depended jointly on maternal and social-male condition, reflecting biparental investment (Courtiol et al., 2016; Kokko & Jennions, 2008). In contrast, fecundity in random and polygynous systems depended only on maternal condition. Under EPCs, offspring paternity was assigned independently to either the social or extra-pair male according to the extra-pair paternity (EPP) share, incorporating cryptic female choice and sperm competition between the social and the extra-pair male (Wan et al., 2013).

Offspring inherit each of 50 loci of the genome-wide paternity via Mendelian segregation (Falconer, 1996)and adaptive traits as mid-parent phenotypic values with mutational deviation, following Porwal (2026). We calculated male reproductive skew according to Wade (1979). The levels of EPP in the simulation runs were taken by rounding off the average and maximum EPP percentage detected in socially monogamous species as reported by Brouwers and Griffith (2019). Simulations were run on the University of Hull “Viper” high-performance computing cluster using the R packages foreach (Microsoft Corporation., 2022a) and doParallel (Microsoft Corporation, 2022b). Each parameter combination was replicated 100 times. Complete parameter ranges, focal values, and formulas used for the calculation of each parameter are provided in Table 1 and Supplementary Tables S1–S2.

**Table 1.** Parameter values assigned for different variables for different mating systems.

| Choice System | $\beta$ Female | $\beta$ Male | Mating System | Parental Care | Female Signal Trait Cost | Male Signal Trait Cost | EPCs | EPC probability | EP P |
| --- | --- | --- | --- | --- | --- | --- | --- | --- | --- |
| **Mutual Choice Monogamy** | 0, 1, 5 | 0, 1, 5 | Monogamy | Equal | 0.25 | 0.25 | TRUE | 0, 10, 30 | 0.2, 0.7 |
| **Random Polygyny** | 0 | 0 | Polygyny | Female only | 0 | 0 | FALSE | 0 | 0 |
| **Female Choice Polygyny** | 0, 1, 5 | 0 | Polygyny | Female only | 0 | 0.25 | FALSE | 0 | 0 |

## Results

Under stable environmental conditions, male reproductive skew and heterozygosity loss varied systematically across mating systems. They were modulated by β (strength of choosiness) and K (carrying capacity). Strict genetic monogamy (EPC = 0) showed the lowest male reproductive skew (close to 0) (Fig. 1) and the slowest loss of heterozygosity (Fig. 2) across all mating scenarios and environmental conditions. However, the difference in reproductive skew between the probability of EPC occurrence (EPC = 10, 30) and female-choice polygynous systems depended on *β*. Higher β increased reproductive skew (Fig. 1) and produced a steeper decline in heterozygosity (Fig. 2) compared with lower *β*. At *β* = 1, EPC = 30 exhibited the highest male reproductive skew (Fig. 1) and, consequently, the steepest decline in heterozygosity over time (Fig. 2). The reproductive skew at *β* = 1 was followed by female-choice polygyny and then by EPC = 10; however, the difference was not evident in terms of heterozygosity loss.

**Figure 1.**
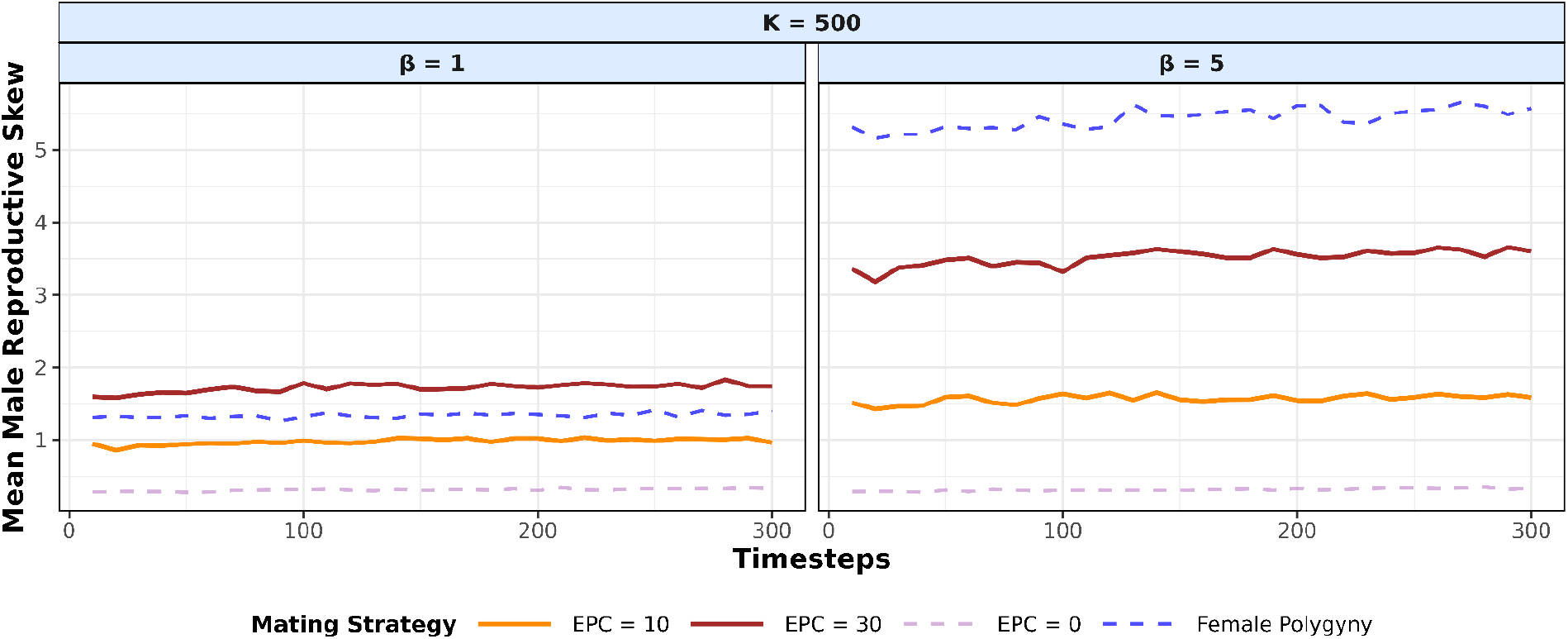
Mean male reproductive skew (y-axis) across 100 replicate populations for each parameter combination over time (x-axis) under stable environmental conditions (i.e., no directional change), homozygosity penalty = 0.5, and *K* = 500, plotted for timesteps where at least 75% of the populations survived. Panels show different levels of choosiness strength (*β*). *β* is applied to both sexes in mutual mate choice scenarios and to females only in female-choice systems. In the EPC mating system, EPP = 0.7.

**Figure 2.**
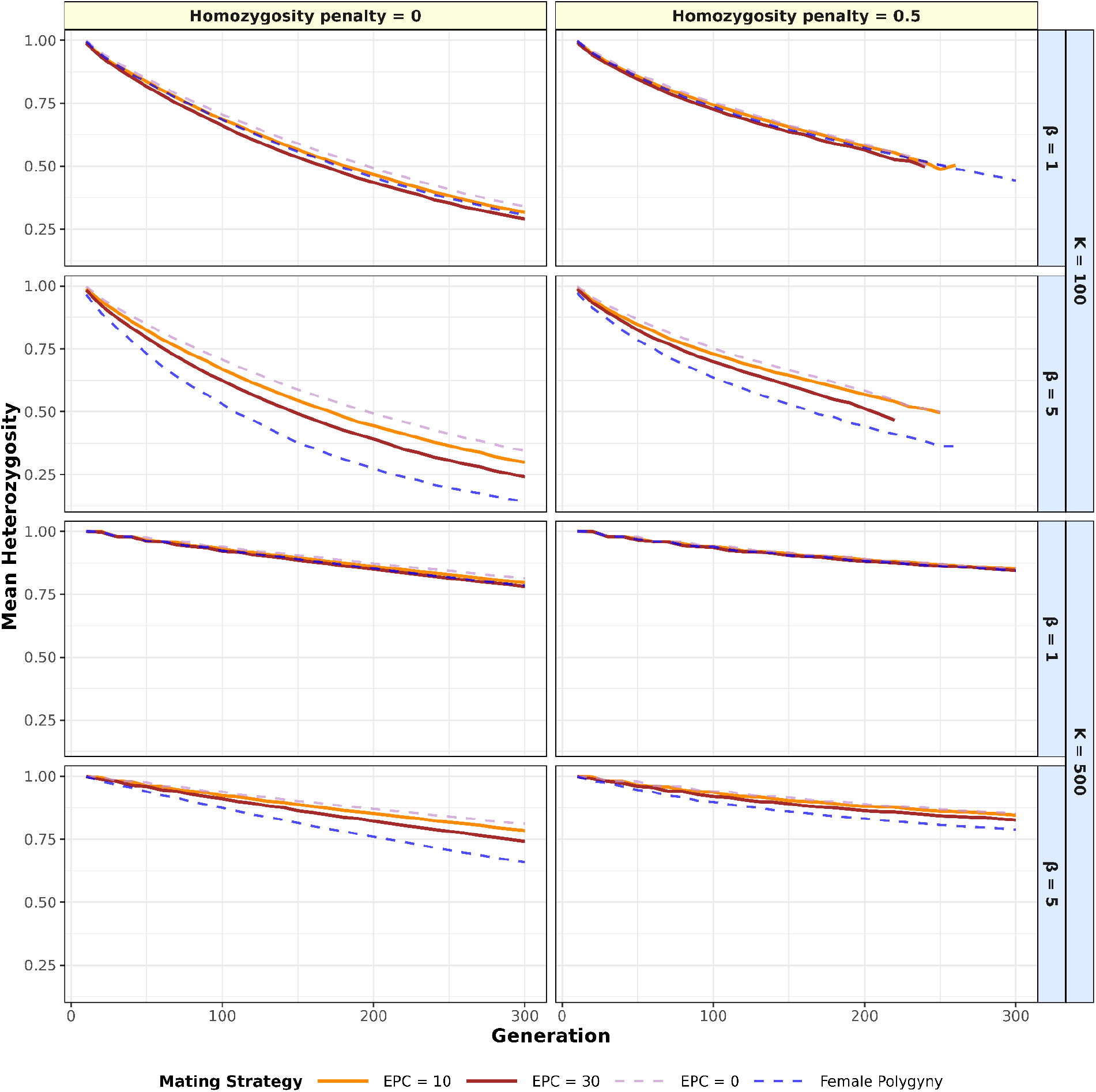
Mean of the median heterozygosity (y-axis) across 100 replicate populations for each parameter combination over time (x-axis) under stable environmental conditions (i.e., no directional change), plotted for timesteps where at least 75% of the populations survived. Vertical panels show different levels of homozygosity penalty, while horizontal panels correspond to varying carrying capacities (*K*) and choosiness strength (*β*). *β* is applied to both sexes in mutual mate choice scenarios and to females only in female-choice systems. In socially monogamous systems where EPC occurs, the rate of extra-pair paternity (EPP) is set to 0.7.

When homozygosity penalties and *β* were elevated at *K* = 100, 75% of the populations went extinct even under stable environmental conditions, with EPC = 30 systems going extinct most rapidly across most replicates (Fig. 2). At *K* = 500, differences among mating systems in the rate of heterozygosity loss became less pronounced, and for all mating systems, at least some populations survived.

Under directional environmental change, population sizes and heterozygosity declined across all mating systems, even during the initial period of relative environmental stability (timesteps 1–50) (Fig. 3, 4, SI1). This decline accelerated as environmental deterioration intensified and the mismatch between the population mean phenotype and the shifting environmental optimum increased. Mutual-choice monogamous systems, with and without EPC, maintained approximately even adult sex ratios, whereas female-choice polygynous systems were consistently female-biased. With more balanced sex ratios, EPC = 0 maintained comparatively higher effective population sizes than systems with EPC = 10, 30, and female-choice polygyny (Fig. 3, 4, SI1). Between social monogamous systems with mutual choice, with EPC = 10, the resultant proportion of EPC out of the total pairs per timestep was between 30 to 50%, whereas for EPC = 30 it was 50 to 80% (Fig. SI1). Female-choice polygynous systems consistently produced more offspring per generation than EPC monogamous systems (Fig. 3, 4).

**Figure 3:**
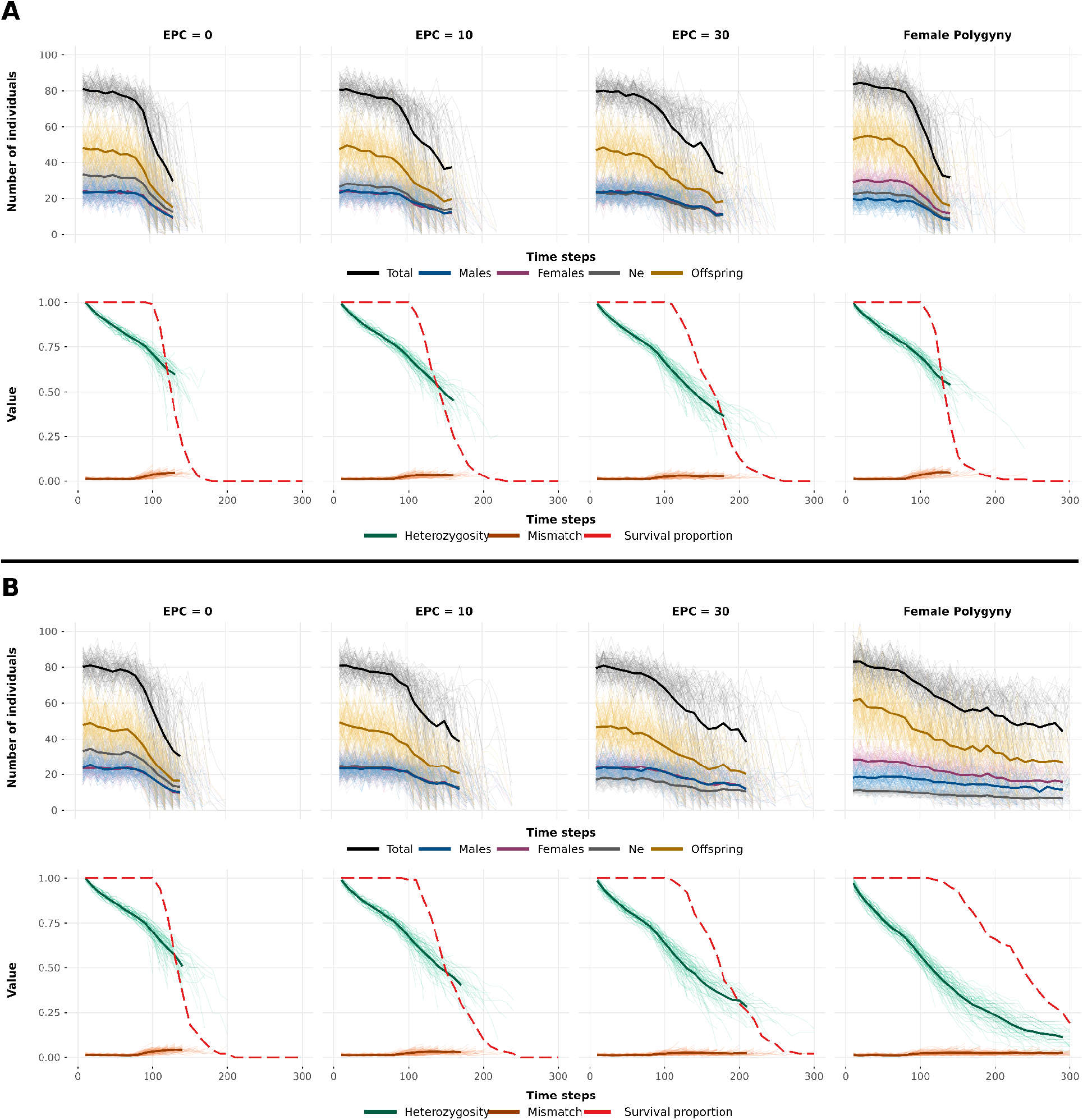
Graphs tracking the means (bold lines) and individual replicates (light lines) of population variables over generation for *K* = 100, homozygosity penalty = 0.25, with A) *β* = 1 and B) *β* = 5 plotted for timesteps where at least 75% of the population survived. In socially monogamous systems where EPC occurs, the rate of extra-pair paternity (EPP) is set to 0.7.

**Figure 4:**
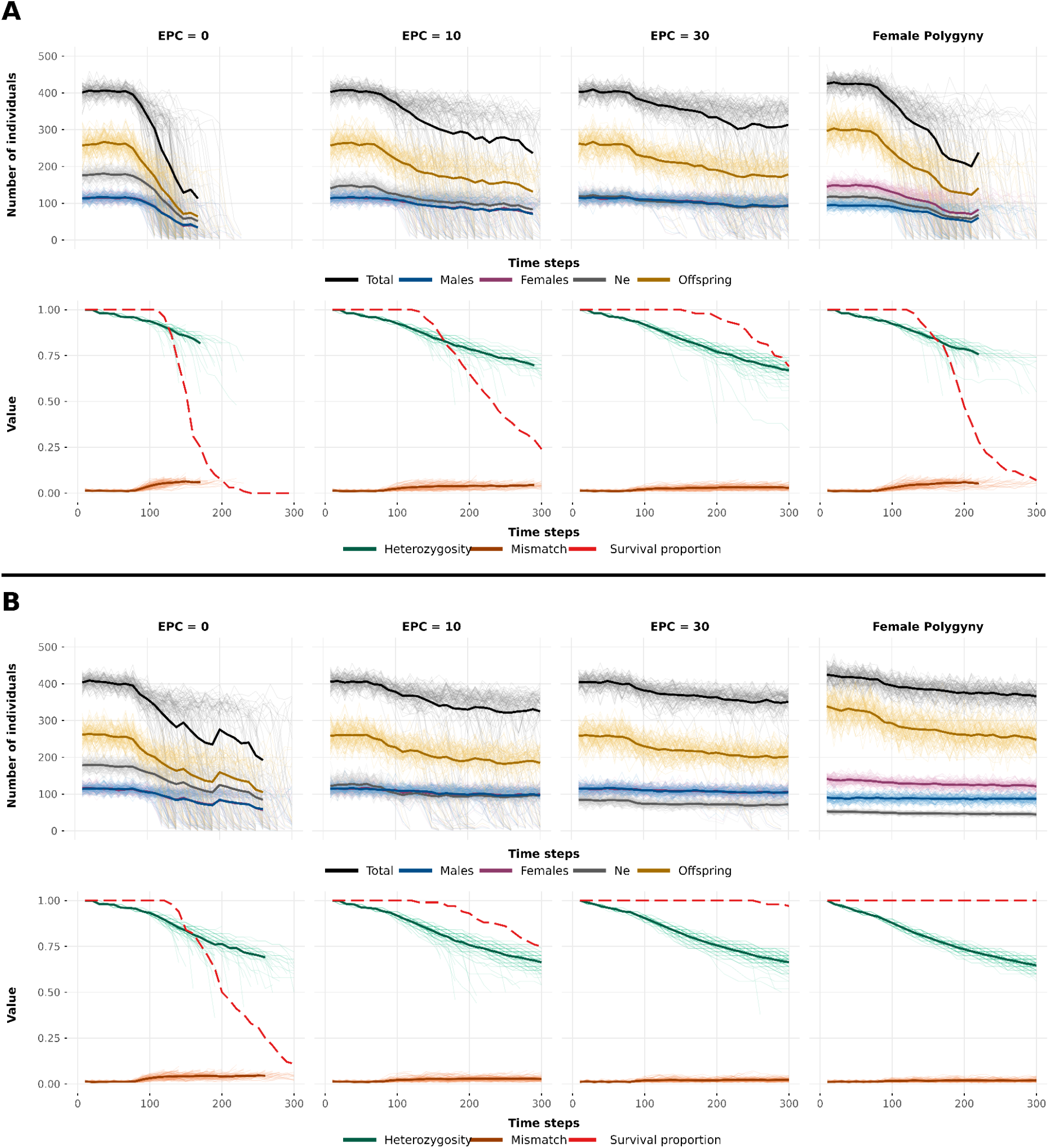
Graphs tracking the means (bold lines) and individual replicates (light lines) over generation for *K* = 500, homozygosity penalty = 0.25, with A) *β* = 1 and B) *β* = 5. In socially monogamous systems where EPC occurs, the rate of extra-pair paternity (EPP) is set to 0.7.

Figure 5 shows that resilience, defined as the rate of directional environmental change at which 50% of replicate populations went extinct, differed substantially among mating systems and was sensitive to homozygosity penalty, EPP, *β*, and *K*. All resilience values are reported relative to the randomly mating polygynous system. Resilience within systems with EPC was sensitive to EPP share and declined as the proportion of offspring sired by the extra-pair male decreased. Under all parameter combinations, EPC = 10 and EPC = 30 outperformed EPC = 0, with EPC = 30 showing the highest resilience among socially monogamous systems. At *β* = 1 with *K* = 100 and homozygosity penalties below 0.25, systems with EPC = 30 showed the highest resilience among all mating systems. Within EPC systems under these conditions, resilience increased as EPP share increased. However, at *K* = 100, when homozygosity penalties exceeded 0.25, resilience fell below that of random polygyny. At the higher carrying capacity (*K* = 500), EPC = 10 and 30 again showed elevated resilience, but only when the extra-pair male sired 80% of all offspring. At *β* = 5, relative resilience increased across all mate-choice-based mating systems compared to lower *β* values. Under this scenario, resilience in EPC = 30 systems was intermediate, consistently higher than in EPC = 0 and 10 but lower than in female-choice polygyny.

**Figure 5.**
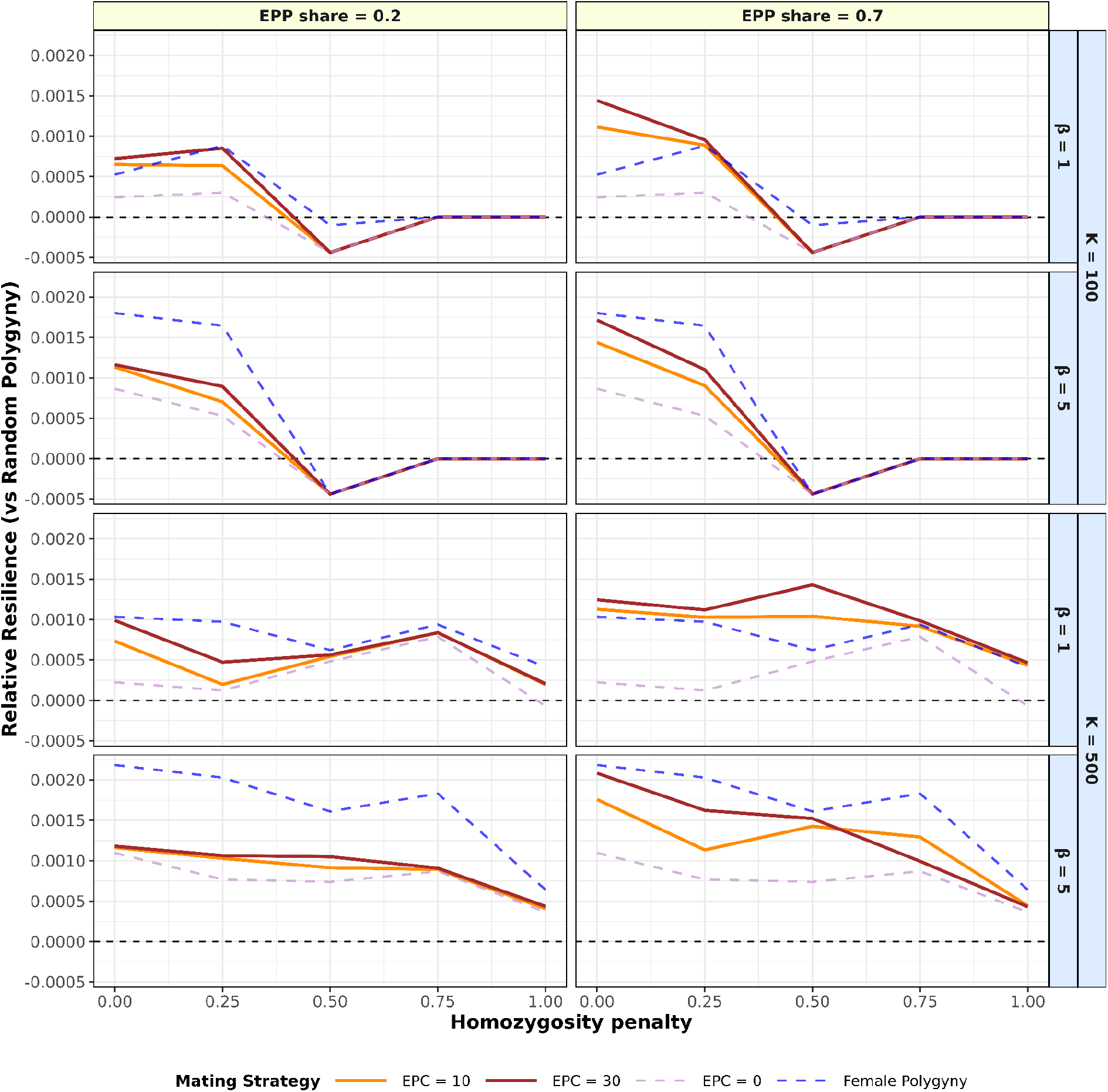
Relative population resilience, defined as the rate of environmental change predicted to drive 50% of populations to extinction, is shown relative to a baseline of randomly mating polygynous systems (resilience - resilience of random polygyny). Vertical panels indicate the proportion of EPP in socially mutual choice monogamous systems. Major horizontal panels represent carrying capacity (*K)*, and nested panels show choosiness strength, applied to both sexes under mutual choice and to females only under female-choice systems. The x-axis shows the strength of homozygosity penalties, and the y-axis shows resilience relative to random polygyny. Lines for mating systems without EPC (EPC=0 and polygyny) are repeated across the EPP panels for comparison. In each panel, the black dashed line denotes the resilience of the random polygynous baseline across levels of homozygosity penalty and K. All parameter combinations were simulated with 100 replicates.

## Discussion

Social monogamy with biparental care is the most prevalent mating system in birds (Cockburn, 2006), but it also occurs in in 3% mammals, including 29% of living primate species (Lukas & Clutton-Brock, 2013), and in some amphibians (Brown et al., 2010; Tumulty et al., 2014). Among birds, a large proportion of species in which social monogamy is accompanied by EPP (Brouer and Griffith, 2019). Still, according to our knowledge, to date extinction risks have only been compared between strict/genetic monogamous and polygamous systems without explicitly modelling varying strength of sexual selection via mate choice (Leach et al., 2020; Legendre et al., 1999; Sæther et al., 2004; Porwal, 2026; Bessa-Gomes et al., 2004). We extended the individual-based model by Porwal et al. (2026) to incorporate and assess how EPC and the degree of EPP (when EPC occurs) in socially monogamous systems with mutual mate-choice shape population resilience to directional environmental change under varying strength of selection based on preference strength. Our results suggest that EPCs play an adaptive role by increasing population resilience relative to socially monogamous systems without EPC as a consequence of females “trading up” for good genes. In some cases, if the likelihood of EPP is high and when the strength of choosiness is low, socially monogamous systems even show higher resilience than female choice polygynous systems under directional environmental change. These context-dependent beneficial effects of EPC are mediated by the level of male reproductive skew generated and depend on the consequences of heterozygosity loss.

Benefits of EPC to population resilience parallel those Powal et al. (2026) reported for polygyny. Although, unlike under polygyny, the total number of offspring in monogamy with EPC remains constrained by the condition of the social male, used here as a proxy for paternal care, thus, EPCs can enhance offspring viability because offspring quality depends on the extra-pair male. By selecting males in better condition during the second round of mating, females effectively shift the system toward a form of female-choice polygyny, promoting a rapid spread of beneficial/adapted genotypes throughout the population. Indeed, our results for EPC and polygyny are similar in that both lead to a decrease in effective population size, as noted by Sugg & Chesser (1994), and a faster loss of heterozygosity compared to monogamy, but increased adaptive rates well compensate for the negative consequences. This process, as a result, enhances long-term population resilience under environmental change. These findings support the hypothesis that EPCs increase adaptive potential under the good genes framework when compared with strict monogamy (Brouwer & Griffith, 2019; Forstmeier et al., 2014; Griffith et al., 2002b; Søraker & Dunning, 2026; Zeh & Zeh, 2001), thereby highlighting the evolutionary advantages of EPCs in changing environments.

However, systems with EPCs are not simply intermediate between monogamy and polygyny in their effects on population resilience. Although socially monogamous systems with mutual mate choice incur notable costs, such as energetic expenses for females expressing signal traits and the exclusion of unpaired individuals due to rigid first-round pair formation, reducing total offspring production (Figs. 3, 4), they can still outperform polygyny when EPP rates are high, provided that both preference strength and the cost of heterozygosity loss are relatively low (Fig. 5). This advantage arises because under high EPP and weak preferences EPC system produces even higher mating skews than polygyny does (Fig. 1), accelerating the spread of adaptive genotypes. When the heterozygosity penalty is low, this advantage is not swamped by the negative effects of heterozygosity loss caused by reproductive bias, resulting in a net positive effect on population resilience compared to polygynous systems.

Our current model thus demonstrates that the superiority of the polygynous system in conferring improved resilience in large populations, as reported in the previous model (Porwal et al. 2026), is not universal. Porwal et al. did not explore the modifying effect of the strength of choosiness, using only *β =5*. Our current model shows that while Porwal et al. (2026)’s conclusion that polygyny is superior to (strict) mutual choice monogamy is robust to higher preference strength, its relative advantage is lower at low *β* when, under certain conditions, polygynous systems may perform worse than the socially monogamous systems with EPC. Thus, the strength of choosiness interacts with mating system in determining extinction risks.

Since lower EPP rates combined with low preference strength do not improve resilience relative to polygyny. This implies that such benefits may not be observed in a substantial proportion of socially monogamous species, given that empirically observed mean EPP occurrence in socially monogamous species is approximately 20% (Brouwer & Griffith, 2019). Finally, while reproductive bias toward high-quality males speeds up adaptation, it simultaneously increases the risk of heterozygosity loss; thus, any resilience benefits associated with high EPC rates remain sensitive to heterozygosity penalties. Nevertheless, such high rates of EPP, even though they have been detected in a few species it may render socially monogamous systems evolutionarily unstable, because the benefits of male parental care are likely to decline with increasing uncertainty of paternity (Søraker et al., 2023), a factor we have not included in our simulations.

Importantly, our model does not account for the costs to females associated with engaging in EPCs, which may constrain female participation in EPC behavior (Søraker & Dunning, 2026). These costs may include harassment from the social male through mate-guarding behaviors (Valera et al., 2003) or coercion and forced copulations by EPC males (McKinney & Evarts, 1998; Wysocki et al., 2025). Future studies should take these costs into account and also consider alternative adaptive hypotheses for EPC occurrence. These include fertility insurance (Sheldon, 1994; Wetton & Parkin, 1991), maximization of genetic compatibility (Kempenaers et al., 1999; Tregenza & Wedell, 2000), and enhancement of genetic diversity (Blomqvist et al., 2002; Foerster et al., 2003; Petrie & Kempenaers, 1998), all of which may be important for predicting population persistence under environmental change.

This model represents an important first step toward understanding the complex dynamics of socially monogamous systems under environmental change. At the same time, it highlights the need for further research on how these systems respond to such conditions in order to design effective conservation strategies. Such strategies may prove critical for protecting a large number of bird populations, particularly those that already exist at low population sizes and are therefore more vulnerable to extinction.

## References

Acerbi, A., Mesoudi, A., & Smolla, M. (2022). Individual-based models of cultural evolution: A step-by-step guide using R. Routledge.

Akçay, E., & Roughgarden, J. (2007). Extra-pair paternity in birds: review of the genetic benefits. Evolutionary ecology research, 9(12), 855–868.

Aquiloni, L., & Gherardi, F. (2008). Mutual mate choice in crayfish: Large body size is selected by both sexes, virginity by males only. Journal of Zoology, 274(2), 171–179. 10.1111/j.1469-7998.2007.00370.x

Arct, A., Drobniak, S. M., & Cichoń, M. (2015). Genetic similarity between mates predicts extrapair paternity—A meta-analysis of bird studies. Behavioral Ecology, 26(4), 959–968.

Baldauf, S. A., Kullmann, H., Schroth, S. H., Thünken, T., & Bakker, T. C. (2009). You can’t always get what you want: Size assortative mating by mutual mate choice as a resolution of sexual conflict. BMC Evolutionary Biology, 9(1), 129. 10.1186/1471-2148-9-129

Barber, C. A., Robertson, R. J., & Boag, P. T. (1996). The high frequency of extra-pair paternity in tree swallows is not an artifact of nestboxes. Behavioral Ecology and Sociobiology, 38(6), 425–430. 10.1007/s002650050260

Beeching, S. C., & Hopp, A. B. (1999). Male mate preference and size-assortative pairing in the convict cichlid. Journal of Fish Biology, 55(5), 1001–1008. 10.1111/j.1095-8649.1999.tb00735.x

Bessa-Gomes, C., Legendre, S., & Clobert, J. (2004). Allee effects, mating systems and the extinction risk in populations with two sexes. Ecology Letters, 7(9), 802–812. 10.1111/j.1461-0248.2004.00632.x

Blomqvist, D., Andersson, M., Küpper, C., Cuthill, I. C., Kis, J., Lanctot, R. B., Sandercock, B. K., Székely, T., Wallander, J., & Kempenaers, B. (2002). Genetic similarity between mates and extra-pair parentage in three species of shorebirds. Nature, 419(6907), 613–615. 10.1038/nature01104

Bolnick, D. I., & Kirkpatrick, M. (2012). The relationship between intraspecific assortative mating and reproductive isolation between divergent populations. Current Zoology, 58(3), 484–492.

Brekke, P., Cassey, P., Ariani, C., & Ewen, J. G. (2013). Evolution of extreme-mating behaviour: Patterns of extrapair paternity in a species with forced extrapair copulation. Behavioral Ecology and Sociobiology, 67(6), 963–972. 10.1007/s00265-013-1522-9

Bro-Jørgensen, J. (2014). Will their armaments be their downfall? Large horn size increases extinction risk in bovids: Large horn size increases extinction risk in bovids. Animal Conservation, 17(1), 80–87. 10.1111/acv.12062

Brook, B. W., Sodhi, N. S., & Bradshaw, C. J. (2008). Synergies among extinction drivers under global change. Trends in Ecology & Evolution, 23(8), 453–460.

Brouwer, L., & Griffith, S. C. (2019). Extra-pair paternity in birds. Molecular Ecology, 28(22), 4864–4882. 10.1111/mec.15259

Brown, J. L., Morales, V., & Summers, K. (2010). A Key Ecological Trait Drove the Evolution of Biparental Care and Monogamy in an Amphibian. The American Naturalist, 175(4), 436–446. 10.1086/650727

Cally, J. G., Stuart-Fox, D., & Holman, L. (2019). Meta-analytic evidence that sexual selection improves population fitness. Nature Communications, 10(1), 2017.

Cockburn, A. (2006). Prevalence of different modes of parental care in birds. Proceedings of the Royal Society B: Biological Sciences, 273(1592), 1375–1383.

Courtiol, A., Etienne, L., Feron, R., Godelle, B., & Rousset, F. (2016). The Evolution of Mutual Mate Choice under Direct Benefits. The American Naturalist, 188(5), 521–538. 10.1086/688658

Daniel, F., Ooi, H., Calaway, R., Microsoft, & Weston, S. (2022). foreach: Provides Foreach Looping Construct (Version 1.5.2) [Computer software]. https://cran.r-project.org/web/packages/foreach/index.html

Falconer, D. S. (1996). Introduction to quantitative genetics. Pearson Education India.

Foerster, K., Delhey, K., Johnsen, A., Lifjeld, J. T., & Kempenaers, B. (2003). Females increase offspring heterozygosity and fitness through extra-pair matings. Nature, 425(6959), 714–717. 10.1038/nature01969

Forstmeier, W., Nakagawa, S., Griffith, S. C., & Kempenaers, B. (2014). Female extra-pair mating: Adaptation or genetic constraint? Trends in Ecology & Evolution, 29(8), 456–464.

Godwin, J. L., Lumley, A. J., Michalczyk, Ł., Martin, O. Y., & Gage, M. J. G. (2020a). Mating patterns influence vulnerability to the extinction vortex. Global Change Biology, 26(8), 4226–4239. 10.1111/gcb.15186

Godwin, J. L., Lumley, A. J., Michalczyk, Ł., Martin, O. Y., & Gage, M. J. G. (2020b). Mating patterns influence vulnerability to the extinction vortex. Global Change Biology, 26(8), 4226–4239. 10.1111/gcb.15186

Griffith, S. C., Owens, I. P. F., & Thuman, K. A. (2002). Extra pair paternity in birds: A review of interspecific variation and adaptive function. Molecular Ecology, 11(11), 2195–2212. 10.1046/j.1365-294X.2002.01613.x

Hasselquist, D., Bensch, S., & von Schantz, T. (1996). Correlation between male song repertoire, extra-pair paternity and offspring survival in the great reed warbler. Nature, 381(6579), 229–232.

Hasselquist, D., & Sherman, P. W. (2001). Social mating systems and extrapair fertilizations in passerine birds. Behavioral Ecology, 12(4), 457–466.

Holveck, M.-J., & Riebel, K. (2010). Low-quality females prefer low-quality males when choosing a mate. Proceedings of the Royal Society B: Biological Sciences, 277(1678), 153–160.

Hooper, P. L., & Miller, G. F. (2008). Mutual Mate Choice Can Drive Costly Signaling Even Under Perfect Monogamy. Adaptive Behavior, 16(1), 53–70. 10.1177/1059712307087283

Jennions, M. D., & Petrie, M. (2000). Why do females mate multiply? A review of the genetic benefits. Biological Reviews, 75(1), 21–64.

Jiang, Y., Bolnick, D. I., & Kirkpatrick, M. (2013). Assortative Mating in Animals. The American Naturalist, 181(6), E125–E138. 10.1086/670160

Johnsen, A., Lifjeld, J. T., Rohde, P. A., Primmer, C. R., & Ellegren, H. (1998). Sexual conflict over fertilizations: Female bluethroats escape male paternity guards. Behavioral Ecology and Sociobiology, 43(6), 401–408.

Jones, I. L., & Hunter, F. M. (1993). Mutual sexual selection in a monogamous seabird. Nature, 362(6417), 238–239. 10.1038/362238a0

Kempenaers, B., Congdon, B., Boag, P., & Robertson, R. J. (1999). Extrapair paternity and egg hatchability in tree swallows: Evidence for the genetic compatibility hypothesis? Behavioral Ecology, 10(3), 304–311. 10.1093/beheco/10.3.304

Kempenaers, B., & Dhondt, A. A. (1993). Why do females engage in extra-pair copulations? A review of hypotheses and their predictions. Belgian Journal of Zoology, 123, 93–93.

Kempenaers, B., Verheyen, G. R., den Broeck, M. V., Burke, T., Broeckhoven, C. V., & Dhondt, A. (1992). Extra-pair paternity results from female preference for high-quality males in the blue tit. Nature, 357(6378), 494–496.

Kempenaers, B., Verheyen, G. R., & Dhondi, A. A. (1997). Extrapair paternity in the blue tit (Parus caeruleus): Female choice, male characteristics, and offspring quality. Behavioral Ecology, 8(5), 481–492.

Kirkpatrick, M., Rand, A. S., & Ryan, M. J. (2006). Mate choice rules in animals. Animal Behaviour, 71(5), 1215–1225.

Kokko, H., & Jennions, M. D. (2008). Parental investment, sexual selection and sex ratios. Journal of Evolutionary Biology, 21(4), 919–948.

Kokko, H., & Johnstone, R. A. (2002). Why is mutual mate choice not the norm? Operational sex ratios, sex roles and the evolution of sexually dimorphic and monomorphic signalling. Philosophical Transactions of the Royal Society of London. Series B: Biological Sciences, 357(1419), 319–330.

Kraak, S. B. M., & Bakker, T. C. M. (1998). Mutual mate choice in sticklebacks: Attractive males choose big females, which lay big eggs. Animal Behaviour, 56(4), 859–866. 10.1006/anbe.1998.0822

Landgraf, C., Wilhelm, K., Wirth, J., Weiss, M., & Kipper, S. (2017). Affairs happen—to whom? A study on extrapair paternity in common nightingales. Current Zoology, 63(4), 421–431.

Lavergne, S., Mouquet, N., Thuiller, W., & Ronce, O. (2010). Biodiversity and Climate Change: Integrating Evolutionary and Ecological Responses of Species and Communities. Annual Review of Ecology, Evolution, and Systematics, 41(1), 321–350. 10.1146/annurev-ecolsys-102209-144628

Leach, D., Shaw, A. K., & Weiss-Lehman, C. (2020). Stochasticity in social structure and mating system drive extinction risk. Ecosphere, 11(2), e03038. 10.1002/ecs2.3038

Lee, A. M., Sæther, B.-E., & Engen, S. (2011). Demographic Stochasticity, Allee Effects, and Extinction: The Influence of Mating System and Sex Ratio. The American Naturalist, 177(3), 301–313. 10.1086/658344

Legendre, S., Clobert, J., Møller, A. P., & Sorci, G. (1999). Demographic Stochasticity and Social Mating System in the Process of Extinction of Small Populations: The Case of Passerines Introduced to New Zealand. The American Naturalist, 153(5), 449–463. 10.1086/303195

Lukas, D., & Clutton-Brock, T. H. (2013). The Evolution of Social Monogamy in Mammals. Science, 341(6145), 526–530. 10.1126/science.1238677

Lumley, A. J., Michalczyk, Ł., Kitson, J. J., Spurgin, L. G., Morrison, C. A., Godwin, J. L., Dickinson, M. E., Martin, O. Y., Emerson, B. C., & Chapman, T. (2015). Sexual selection protects against extinction. Nature, 522(7557), 470–473.

Martínez-Ruiz, C., & Knell, R. J. (2017). Sexual selection can both increase and decrease extinction probability: Reconciling demographic and evolutionary factors. Journal of Animal Ecology, 86(1), 117–127. 10.1111/1365-2656.12601

Martinossi-Allibert, I., Savković, U., \DJor\djević, M., Arnqvist, G., Stojković, B., & Berger, D. (2018).The consequences of sexual selection in well-adapted and maladapted populations of bean beetles. Evolution, 72(3), 518–530.

Martins, M. J. F., Puckett, T. M., Lockwood, R., Swaddle, J. P., & Hunt, G. (2018). High male sexual investment as a driver of extinction in fossil ostracods. Nature, 556(7701), 366–369.

McKaye, K. R. (1986). Mate choice and size assortative pairing by the cichlid fishes of Lake Jiloá, Nicaragua. Journal of Fish Biology, 29(sA), 135–150. 10.1111/j.1095-8649.1986.tb05005.x

McKinney, F., & Evarts, S. (1998). Sexual Coercion in Waterfowl and Other Birds. Ornithological Monographs, (49), 163–195. 10.2307/40166723

Mennill, D. J., Boag, P. T., & Ratcliffe, L. M. (2003). The reproductive choices of eavesdropping female black-capped chickadees, Poecile atricapillus. Naturwissenschaften, 90(12), 577–582. 10.1007/s00114-003-0479-3

Møller, A. P., & Birkhead, T. R. (1994). THE EVOLUTION OF PLUMAGE BRIGHTNESS IN BIRDS IS RELATED TO EXTRAPAIR PATERNITY. Evolution, 48(4), 1089–1100. 10.1111/j.1558-5646.1994.tb05296.x

Moore, M. P., Nalley, S. E., & Hamadah, D. (2024). An evolutionary innovation for mating facilitates ecological niche expansion and buffers species against climate change. Proceedings of the National Academy of Sciences, 121(10), e2313371121. 10.1073/pnas.2313371121

Morrow, E. H., & Fricke, C. (2004). Sexual selection and the risk of extinction in mammals. Proceedings of the Royal Society of London. Series B: Biological Sciences, 271(1555), 2395–2401.

Morrow, E. H., & Pitcher, T. E. (2003). Sexual selection and the risk of extinction in birds. Proceedings of the Royal Society of London. Series B: Biological Sciences, 270(1526), 1793–1799.

Otter, K., Ratcliffe, L., Michaud, D., & Boag, P. T. (1998). Do female black-capped chickadees prefer high-ranking males as extra-pair partners? Behavioral Ecology and Sociobiology, 43(1), 25–36.

Parrett, J. M., Mann, D. J., Chung, A. Y. C., Slade, E. M., & Knell, R. J. (2019). Sexual selection predicts the persistence of populations within altered environments. Ecology Letters, 22(10), 1629–1637. 10.1111/ele.13358

Petrie, M., & Kempenaers, B. (1998). Extra-pair paternity in birds: Explaining variation between species and populations. Trends in Ecology & Evolution, 13(2), 52–58. 10.1016/S0169-5347(97)01232-9

Petrie, M., Tim, H., & Carolyn, S. (1991). Peahens prefer peacocks with elaborate trains. Animal Behaviour, 41(2), 323–331.

Rätti, O., Hovi, M., Lundberg, A., Tegelström, H., & Alatalo, R. V. (1995). Extra-pair paternity and male characteristics in the pied flycatcher. Behavioral Ecology and Sociobiology, 37(6), 419–425.

Román-Palacios, C., & Wiens, J. J. (2020). Recent responses to climate change reveal the drivers of species extinction and survival. Proceedings of the National Academy of Sciences, 117(8), 4211–4217. 10.1073/pnas.1913007117

Rueger, T., Gardiner, N. M., & Jones, G. P. (2016). Size matters: Male and female mate choice leads to size-assortative pairing in a coral reef cardinalfish. Behavioral Ecology, arw082.

Sæther, B.-E., Engen, S., Lande, R., Møller, A. P., Bensch, S., Hasselquist, D., Beier, J., & Leisler, B. (2004). Time to extinction in relation to mating system and type of density regulation in populations with two sexes. Journal of Animal Ecology, 925–934.

Saino, N., Primmer, C. R., Ellegren, H., & M⊘ller, A. P. (1997). An experimental study of paternity and tail ornamentation in the barn swallow (Hirundo rustica). Evolution, 51(2), 562–570.

Salamin, N., Wüest, R. O., Lavergne, S., Thuiller, W., & Pearman, P. B. (2010). Assessing rapid evolution in a changing environment. Trends in Ecology & Evolution, 25(12), 692–698. 10.1016/j.tree.2010.09.009

Sheldon, Ben. C. (1994). Male phenotype, fertility, and the pursuit of extra-pair copulations by female birds. Proceedings of the Royal Society B: Biological Sciences, 257(1348), 25–30. 10.1098/rspb.1994.0089

Søraker, J. S., & Dunning, J. (n.d.). The costs of extra-pair behaviours in birds. Biological Reviews, n/a(n/a). 10.1002/brv.70178

Søraker, J. S., Wright, J., Hanslin, F. Ø., & Pepke, M. L. (2023). The evolution of extra-pair paternity and paternal care in birds. Behavioral Ecology, 34(5), 780–789. 10.1093/beheco/arad053

Sorci, G., Møller, A. P., & Clobert, J. (1998). Plumage dichromatism of birds predicts introduction success in New Zealand. Journal of Animal Ecology, 67(2), 263–269. 10.1046/j.1365-2656.1998.00199.x

Strohbach, S., Curio, E., Bathen, A., Epplen, J., & Lubjuhn, T. (1998). Extrapair paternity in the Great Tit (Parus major): A test of the “good genes” hypothesis. Behavioral Ecology, 9(4).

Sugg, D. W., & Chesser, R. K. (1994). Effective Population Sizes with Multiple Paternity. Genetics, 137(4), 1147–1155. 10.1093/genetics/137.4.1147

Team, R. C. (2023). R: The R project for statistical computing.

Tregenza, T., & Wedell, N. (2000). Genetic compatibility, mate choice and patterns of parentage: Invited Review. Molecular Ecology, 9(8), 1013–1027. 10.1046/j.1365-294x.2000.00964.x

Tumulty, J., Morales, V., & Summers, K. (2014). The biparental care hypothesis for the evolution of monogamy: Experimental evidence in an amphibian. Behavioral Ecology, 25(2), 262–270.

Urban, M. C. (2024). Climate change extinctions. Science, 386(6726), 1123–1128. 10.1126/science.adp4461

Valera, F., Hoi, H., & Krištín, A. (2003). Male shrikes punish unfaithful females. Behavioral Ecology, 14(3), 403–408. 10.1093/beheco/14.3.403

Wade, M. J. (1979). Sexual Selection and Variance in Reproductive Success. The American Naturalist, 114(5), 742–747. 10.1086/283520

Wan, D., Chang, P., & Yin, J. (2013). Causes of extra-pair paternity and its inter-specific variation in socially monogamous birds. Acta Ecologica Sinica, 33(3), 158–166. 10.1016/j.chnaes.2013.03.006

Weston, S. (2022). Microsoft corporation. doParallel: Foreach parallel adaptor for the ‘parallel’package. R package version 1.0. 17.

Wetton, J. H., & Parkin, D. T. (1991). An association between fertility and cuckoldry in the house sparrow, Passer domesticus. Proceedings of the Royal Society B: Biological Sciences, 245(1314), 227–233. 10.1098/rspb.1991.0114

Whittingham, L. A., & Dunn, P. O. (2001). Survival of extrapair and within-pair young in tree swallows. Behavioral Ecology, 12(4), 496–500.

Whittingham, L. A., & Dunn, P. O. (2016). Experimental evidence that brighter males sire more extra-pair young in tree swallows. Molecular Ecology, 25(15), 3706–3715. 10.1111/mec.13665

Wysocki, D., Cholewa, M., & Halupka, K. (2025). Biological significance of forced extra-pair copulations in a population of European Blackbirds. Scientific Reports, 15(1), 8974. 10.1038/s41598-025-92997-4

Zeh, J. A., & Zeh, D. W. (2001). Reproductive mode and the genetic benefits of polyandry. Animal Behaviour, 61(6), 1051–1063. 10.1006/anbe.2000.1705

