## Supplementary Information for "Population Resilience Under Environmental Deterioration in Socially Monogamous Systems with Mutual Mate Choice"

**Table S1**: State Variables and Parameters

| **Parameter/Variable** | **Description** | **Type** | **Values/Range** |
| --- | --- | --- | --- |
| **Population Variables** |  |  |  |
| b | Probability of birth per offspring attempt | Fixed | 0.5 |
| K | Carrying capacity of the population | Fixed | 100, 500 |
| t | Total number of time steps in the simulation | Fixed | 200 |
| e | Initial environmental value | Variable | 0.2 |
| mating_system | Type of mating system | Fixed | "monogamy" or "polygyny" |
| epc | Extra-pair copulation (EPC) allowed | Fixed | TRUE/FALSE |
| epc_prob | Strength of EPC tendency | Fixed | 10, 30 |
| care weight male | Determines if offspring survival depends on mismatch and heterozygosity of both parents’ (0.5) or only mother’s (0) | Fixed | 0.5 (for mutual mate choice monogamy) or 0 (for rest of the mating systems) |
| Extra pair paternity (EPP) | Fraction of offspring sired by the EPC male during EPC | Fixed | 0, 0.2, 0.8 |
| **Individual Variables** |  |  |  |
| O | Maximum number of offspring per female | Fixed | 5 |
| I | Age of sexual maturity | Fixed | 2 |
| grp.m | Number of males assessed by females per mating group | Fixed | 10 |
| grp.f | Number of females assessed by males per mating group | Fixed | 10 |
| alpha.m (α_m)_ | Strength of condition dependence for male sexual trait | Fixed | 2 |
| alpha.f (α*f*) | Strength of condition dependence for female sexual trait | Fixed | 2 |
| cost.m | Cost of sexual trait expression in males affecting survival | Fixed | 0, 0.25 |
| cost.f | Cost of sexual trait expression in females affecting survival | Fixed | 0, 0.25 |
| beta.m (*β*) | Male mate preference strength | Fixed | 0, 1, 5 |
| beta.f (*β*) | Female mate preference strength | Fixed | 0, 1, 5 |
| homozygosity penalty | Scaling factor penalizing low heterozygosity in survival, reproduction and signal trait | Fixed | 0, 0.25, 0.5, 0.75, 1 |
| mismatch.penalty | Scaling factor penalizing mismatch in survival, reproduction and signal trait | Fixed | 5 |
| alive | Individual survival status | Variable | 1 (alive) or 0 (dead) |
| sex | Individual sex | Fixed | "M" (male) or "F" (female) |
| age | Individual age | Variable | 0 to 10 |
| heterozygosity (*H*) | Proportion of loci where genetic strands differ | Variable | 0 to 1 |
| phenotype | Individual quantitative phenotypic trait value determining adaptation | Variable | Normal distribution, mean = 0.2 (initial) |
| Environmental Variables |  |  |  |
| e.var | Mode of environmental variation | Fixed | “Directional" |
| directional.rate | Mean change per time step | Fixed | 0 (stable environment), 0.001, 0.002, 0.003, 0.004 |
| **Genetic Variables** |  |  |  |
| loci | Number of genetic loci for heterozygosity calculation in each strand | Fixed | 50 (default) |
| mutation_phe | Standard deviation of phenotypic mutation | Fixed | 0.01 (default) |

**Table S2**: Submodels and Mathematical Formulations

| Submodel | Description | Mathematical Formulation |
| --- | --- | --- |
| Environmental Variation | Updates the environmental value 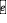based on the mode of variation. | For e.var = "Directional":  - If directional.rate = 0, 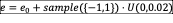  Else if 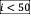, 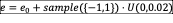  otherwise, 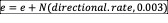 |
| Mismatch (*M*) | Computes mismatch as the absolute difference between phenotype and environment. | *\|e – phenotype\|* |
| Signal Trait | Calculates sex-specific signal trait for mature individuals (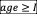). | For 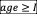:  Males: 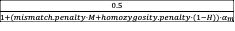-  Females: For 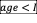: 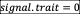 |
| Death Probability | Determines survival probability based on density, mismatch, heterozygosity, and signal trait costs. | 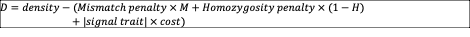  density = *K* / population size, mismatch penalty and inbreeding penalty are variable, and cost (*cost.m* for males, *cost.f* for females,) represents the survival cost of expressing the signal trait. |
| Mate Choice | Assigns mating pairs based on signal traits, preference strengths, and mating system. | - Group assignment:  Females to 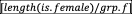 groups;  males to 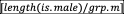 groups.    Female choice probability: If 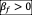 and group size 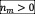,  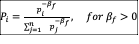, where 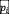is male rank; else 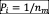.    Male choice probability: if 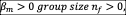   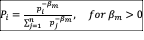, where 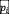is female rank; else 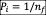. |
| Probability of EPC occurrence | Determines the likelihood of a female mating again with with a male other than the social male | 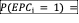  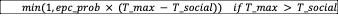  *epc_prob = probability scaling factor for EPC*  *T_max = highest male signal trait in the mating group*  *T_social = signal trait of the female’s social mate* |
| Reproduction | Determines number of offspring based on parental mismatch and heterozygosity. | 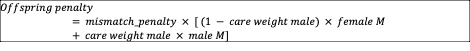  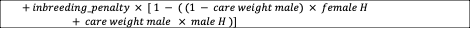  Number of offspring:  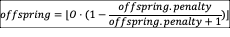 |
| Effective Population Size | Calculates effective population size based on variance in reproductive success. | Female effective size:  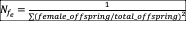  Male effective size: 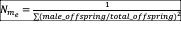  Effective population size: 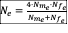if 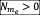 and 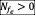, else 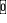. |
| Male reproductive skew | Based on male reproductive success | Male reproductive skew = variance in offspring sired across all eligible males / mean number of offspring sired per eligible male |

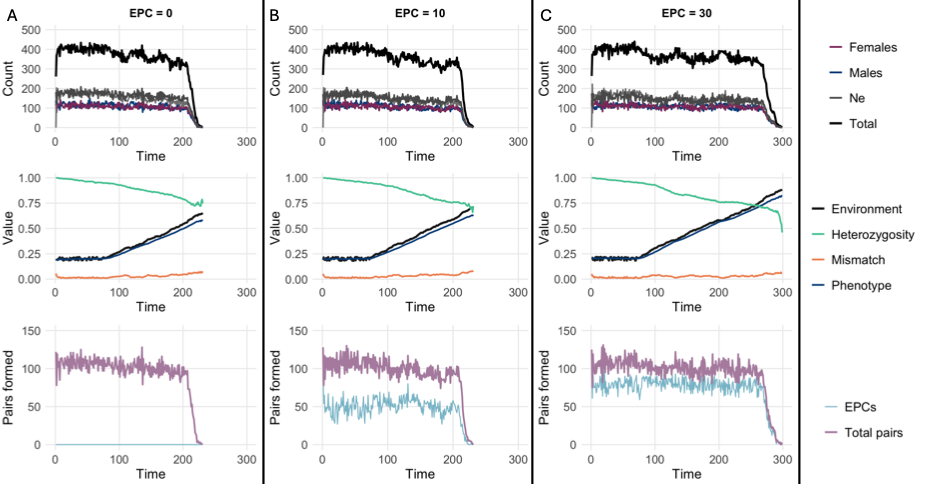

**Figure SI1.** Representative outputs from simulation runs under three mating systems: (A) EPC = 0, (B) EPC = 30, and (C) female-choice polygyny. Simulations were conducted with a directional change rate of 0.003, mate preference strength (*β*) = 0.5, and a homozygosity penalty of 0.5. The top row of panels shows population demography over time. The middle row depicts environmental conditions and the population’s trait values relative to the environment. The bottom row illustrates the total number of pairings formed and, where applicable, the number of EPC events (with EPP = 0.8 in EPC scenarios).
